# Synergistic activation of RARb and RARg nuclear receptors restores cell specialization during stem cell differentiation by hijacking RARa-controlled programs

**DOI:** 10.1101/2022.05.31.494116

**Authors:** Aysis Koshy, Elodie Mathieux, François Stüder, Aude Bramoulle, Michele Lieb, Bruno Maria Colombo, Hinrich Gronemeyer, Marco Antonio Mendoza-Parra

**Affiliations:** UMR 8030 Génomique Métabolique, Genoscope, Institut François Jacob, CEA, CNRS, University of Evry-val-d’Essonne, University Paris-Saclay, 91057 Évry, France; Department of Functional Genomics and Cancer, Institut de Génétique et de Biologie Moléculaire et Cellulaire, Illkirch, France

**Keywords:** retinoids, cell specialization, gene programs, neuronal differentiation

## Abstract

How cells respond to different external cues to develop along defined cell lineages to form complex tissues is a major question in systems biology. Here, we investigated the potential of retinoic acid receptor (RARs)-selective synthetic agonists to activate the gene-regulatory programs driving cell specialization during nervous tissue formation from P19 stem cells. Specifically, we found that the synergistic activation of the RARβ and RARγ by selective ligands (BMS641 or BMS961) induces cell maturation to specialized neuronal subtypes, as well as to astrocytes and oligodendrocyte precursors.

Using RAR istoype knockout lines exposed to RAR-specific agonists, interrogated by global transcriptome landscaping and *in silico* modeling of transcription regulatory signal propagation, revealed major RARα−driven gene programs essential for optimal neuronal cell specialization, and hijacked by the synergistic activation of the RARβ and RARγ receptors.

Overall, this study provides a systems biology view of the gene programs accounting for the previously observed redundancy between RAR receptors, paving the way towards their potential use for directing cell specialization during nervous tissue formation.

## Introduction

The potential of all-*trans* retinoid acid (ATRA) to induce differentiation of embryonic stem and embryonic carcinoma (EC) cells is well established (Dollé and Niederreither, 2015; Soprano et al., 2007). ATRA is a ligand for the three retinoic acid receptors (RARα, β and γ) and major medicinal chemistry efforts have resulted in the synthesis of ligands that are selective for each RAR isotype (Álvarez *et al*, 2014; de Lera *et al*, 2007). Multiple studies, including ours, demonstrated that P19 stem cells differentiate into neuronal precursors when treated with ATRA or the RARα-specific agonist BMS753, but they do not progress in differentiation when treated with the RARβ-specific agonist BMS641 or the RARγ-specific agonist BMS961 (Mendoza-Parra *et al*, 2016a).

Here we have investigated the neuronal lineage-inducing potential of individual and combined subtype-specific retinoids in 2-dimensional monolayer culture. We observe that, in addition to ATRA and RARα agonists, the combination of RARβ and RARγ agonists triggers a complex differentiation process generating a variety of neuronal subtypes as well as oligodendrocyte precursors and GFAP (+) astrocytes. This synergistic effect has been decorticated on the grounds of the RAR/RXR-driven gene programs, and the use of RAR subtype-deficient cells, which were instrumental for revealing the specificity of each of the synthetic ligands. Finally, we reveal that the RARβ+γ synergy, which involves a defined set of gene programs controlled by key master players, is antagonized in the presence of RARα, suggesting that an asynchronous activation of the various RAR receptors leads to impaired neuronal specialization.

## Results

### Synergistic activation of RARγ and RARβ induces neuronal cell specialization in P19 embryonic stem cells

Using the well-established monolayer culture for efficient morphological P19 cell differentiation (Monzo *et al*, 2012)(Mendoza-Parra et al., 2016), we observed that after 10 days of treatment, ATRA or the RARα agonist BMS753 induced neuronal precursors, as revealed by immunofluorescence using the neuronal marker tubulin β-3 (TUBB3) but also mature neurons, as revealed by the microtubule-associated protein 2 (MAP2) (**Figure 1A&B**). In contrast, treatment with the RARβ-specific ligand BMS641 or the RARγ agonist BMS961 did not lead to neuronal differentiation. We only observed neuronal-like cells presenting short neurite outgrowth structures, devoid of MAP2 immunostaining. Surprisingly, the combination of these two synthetic agonists (BMS641 & BMS961) restored neuronal differentiation presenting neurite outgrowth characteristics as similar as those observed on ATRA or BMS753 treated samples (**Figure 1B**).

**Figure 1.**
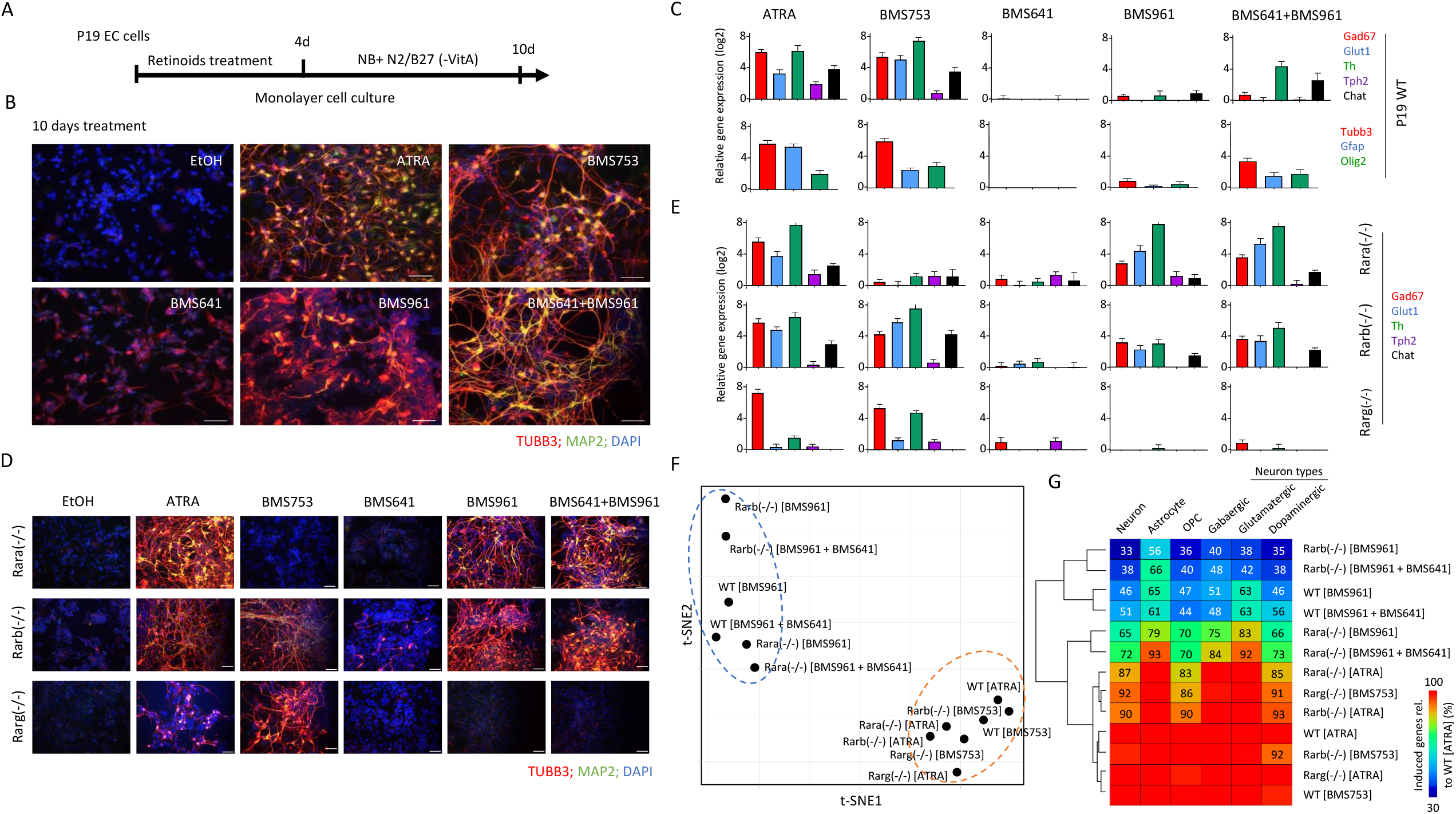
Synergistic activation of RARγ and RARβ induces neuronal cell specialization in P19 embryonic stem cells. **(A)** Schematic representation of the P19 cell differentiation assay. P19 cells cultured on monolayer are exposed to retinoids during 4 days to induce cell fate commitment; then they are cultured during 6 more days on a synthetic medium (Neurobasal: NB) complemented with N2 and B27(without Vitamin A) supplements. **(B)** Immuno-fluorescence micrograph of wild-type P19 cells after 10 days of culture in presence of either Ethanol (EtOH: vehicle control), all-trans retinoic acid (ATRA), the RARa agonist BMS753, the RARb agonist BSM641, the RARg agonist BMS961 or the combination of RARb and RARg agonists. Cells were stained for the neuronal precursor marker TUBB3 (red) and the marker for mature neurons MAP2 (green). Nuclei were stained with DAPI (blue). **(C)** Top panel: RT-qPCR revealing the mRNA expression levels of gene markers associated to GABAergic (Gad67), glutamatergic (Glut1), dopaminergic (Th), or cholinergic (Chat) neuronal subtypes in samples treated with the indicated RAR agonists. Bottom panel: RT-qPCR mRNA expression levels of the glial fibrillary acidic proteins (Gfap) and oligodendrocyte transcription factor 2 (Olig2) genes. **(D)** Immuno-fluorescence micrograph of P19 Rar-null mutant cells after 10 days of treatment with the aforementioned RAR agonists. **(E)** RT-qPCR mRNA expression levels of gene markers associated to the aforementioned neuronal subtypes assessed on P19 Rar-null mutant cells. **(F)** T-distributed stochastic neighbor embedding (t-SNE) analysis of differential gene expression readouts assessed on global transcriptomes performed on WT or RAR-null cells treated with specific agonists (10 days). Differential gene expression has been assessed realtive to the Ethanol treated control sample (fold change levels >4). **(G)** Fraction of upregulated genes (fold change levels >4) associated to markers corresponding to specialized cells relative to those observed on the gold-standard wild-type ATRA treated sample. Fraction levels higher than 95% are only displayed with the heatmap color-code (red).

Neuronal maturation has been further supported by RT-qPCR assays revealing significant transcript levels associated to markers for GABAergic (Gad67), glutamatergic (Glut1), dopaminergic (Th), or cholinergic (Chat) neuronal subtypes in samples treated with ATRA and BMS753 (**Figure 1C**). Combined exposure to the RARβ+γ agonists (BMS641, BMS961) presented significant expression levels only for the markers Th and Chat, suggesting a partial neuronal subtype differentiation in comparison to ATRA or BMS753 treatment, but also the necessity of a more comprehensive strategy (global transcriptomes) to evaluate the cell specialisation success. In addition to neuronal cell specialization, RT-qPCR assays also revealed significant expression levels of the glial fibrillary acidic proteins (Gfap) and oligodendrocyte transcription factor 2 (Olig2) genes, indicative for the presence of astrocyte and oligodendrocyte precursors, both in ATRA and BMS753 treatment, as well as in the combination of the BMS641 & BMS961 agonists (**Figure 1C**).

While the combined exposure to the RARβ+γ agonists (BMS641, BMS961) led to morphological neuronal cell specialization, the evaluated markers present systematic lower levels than those observed in ATRA or BMS753, As this could be due to a potential inhibitory effect of non-liganded RARα, we engineered P19 cells deficient for each of the RARs using the CRISPR/Cas9 technology. Surprisingly, the absence of expression of either RARα, RARβ or RARγ receptor directly affected expression of the non-deleted RARα and the RARβ receptors, notably by preserving their induction after 96 hours of treatment (**Suppl. Figure S1**). Furthermore, P19 Rara(-/-) cells gave rise to mature neurons when treated with ATRA, but also with the RARγ agonist BMS961 or the combination of RARβ+γ ligands (BMS641 & BMS961), as revealed by TUBB3/MAP2 immunostaining (**Figure 1D**) and the high expression of transcripts associated to neuronal subtypes Gad67, Glut1 and Th (**Figure 1E**), as well as Tubb3 and the oligodendrocyte marker Olig2 (**Suppl. Figure S2**). The observed enhanced neuronal differentiation in presence of the RARγ agonist in P19 Rara(-/-) relative to WT cells is in agreement with previous studies on the functional redundancy of RAR subtypes during endodermal (F9) and neuronal (P19) cell differentiation (Taneja *et al*, 1996, 1995; Roy *et al*, 1995). Neuronal differentiation was also observed in RARβ-deficient cells treated with ATRA or the RARα agonist (BMS753), but also the RARγ agonist (BMS961) and the RARβ+γ agonist-combination (BMS641 & BMS961) (**Figure 1D&E**). Finally, the Rarg(-/-) cells entered neuronal differentiation only in presence of ATRA or BMS753, in agreement with our earlier finding that the RARα-dependent gene program directs the neuronal cell fate of P19 cells (Mendoza-Parra *et al*, 2016a).

While neuronal differentiation performance driven by RARβ+γ agonist treatment on WT, Rara- and Rarb-deficient cells, was evaluated by immunostaining and RT-qPCR assays targeting few marker genes, we reasoned that a comprehensive strategy could reveal potential differences among these multiple conditions. Global transcriptome assays performed on WT or RAR subtype-deficient cells treated with specific agonists during 10 days revealed between 1,340 and 2,250 up-regulated genes (fold change levels >4 relative to the ethanol control) allowing to query for cell specialization signatures and their corresponding divergencies between samples (**Supp. Figure S3**). Indeed, a t-distributed stochastic neighbor embedding (t-SNE) analysis of their differentially expressed genes revealed two major groups. The first comprises WT or RAR subtype-deficient cells treated with either ATRA or the RARa agonist (BMS753) (**Figure 1F**). The second group gathers samples treated with the RARγ (BMS961) or the combination of the RARβ and RARγ agonists (BMS641 & BMS961). This second group displays significant disparities among their components, with the transcriptomes of RARβ+γ-agonist-treated Rara(-/-) cells being closer to the group 1 than the others, in line with the observed neuronal differentiation (**Figure 1 F**).

To further understand this classification through a better characterization of the cell specialization signature during these differentiation conditions, we have collected an ensemble of gene markers associated to neurons (1,352 genes), astrocytes (501 genes), oligodendrocyte precursors (OPC: 501 genes) as well as on GABAergic (318 genes), glutamatergic (311 genes) and dopaminergic (513 genes) neuronal subtypes (**Suppl. Table S1**) (Tasic *et al*, 2018; Hook *et al*, 2018; Voskuhl *et al*, 2019). By comparing the number of upregulated genes in the WT ATRA treatment with these comprehensive lists of markers, we have revealed that ∼30% of them are associated to markers corresponding to specialized cells (632 from 2,158 upregulated genes from which 401 are associated to neurons, 100 to astrocytes and 131 to oligodendrocyte precursors) (**Suppl. Figure S3**). Considering the upregulated genes associated to specialized cells in the WT ATRA condition as gold-standard for optimal cell differentiation, we have revealed that most of the ATRA or BMS753-treated samples presented similar amounts of genes associated to specialized cells. Indeed, the Rara(-/-) mutant sample treated with ATRA recapitulated ∼83% of the gold-standard up-reguated genes associated to oligodendrocyte precursors, ∼87% for neuronal associated gene markers, and ∼85% for the dopaminergic neuron subtype; and a similar behavior is observed for the Rarb(-/-) mutant treated with ATRA or Rarg(-/-) mutant the treated with the BMS753 ligand (**Figure 1 G**). In contrary, samples treated with the RARγ agonists BMS961 give rise to cell marker levels of only ∼30% in the context of the Rarb(-/-) mutant, to ∼70% for the Rara(-/-). Importantly, treating the Rara(-/-) mutant with the combination of the RARβ+γ-agonist (BMS641 BMS961) leads to >70% of the gold-standard levels associated to neuronal cells, and even more than 90% for astrocyte or the glutamatergic neuronal cell type, demonstrating that the use of a synergistic RARβ+γ-agonist-treatment on RARa subtype-deficient cells leads to an enhanced restoration of cell specialization during P19 stem cell differentiation.

### P19 differentiation driven by the combination of RARβ and RARγ agonists present a delayed expression of cell specialization markers

To assess the temporal evolution of gene expression during RAR ligand-induced neuronal differentiation and to associate specific gene programs with the appearance of neuronal cell subtypes, we generated global transcriptomes after 2, 4 and 10 days of treatment of WT P19 cells.This has been performed for samples treated with either the pan agonist ATRA – as a gold-standard treatment –, the RARα specific agonist BMS753 or the combination of the RARβ and RARγ agonists (BMS641 & BMS961), shown to induce neuronal cell specialization.

Differential gene expression through the aforementioned time-points - assessed during ATRA treatment - were classified on 14 relevant gene co-expression paths, defined herein as a group of genes with similar temporal changes of expression levels (**Figure 2A**). For instance, Path1 (1,195 genes) corresponds to genes upregulated (fold-change >2 relative to vehicle at d0) after two days of treatment and remained over-expressed till day 10. Paths 2 (725 genes) and 4 (582 genes) comprise late upregulated genes, induced only at d4 or d10, respectively (**Figure 2A**). A gene ontology term analysis performed over the first seven gene co-expression paths – associated to up-regulated genes at least at one time-point – reveals that Path1 comprises genes involved in neuron differentiation, nervous system development or axon guidance; Path 2 in axonogenesis or axon development; while Path4 is associated with chemical synaptic transmission, synaptic vesicle budding or synapse organization; in agreement with the time of induction along the neuronal differentiation lineage and sub-type specification (**Figure 2B**).

**Figure 2.**
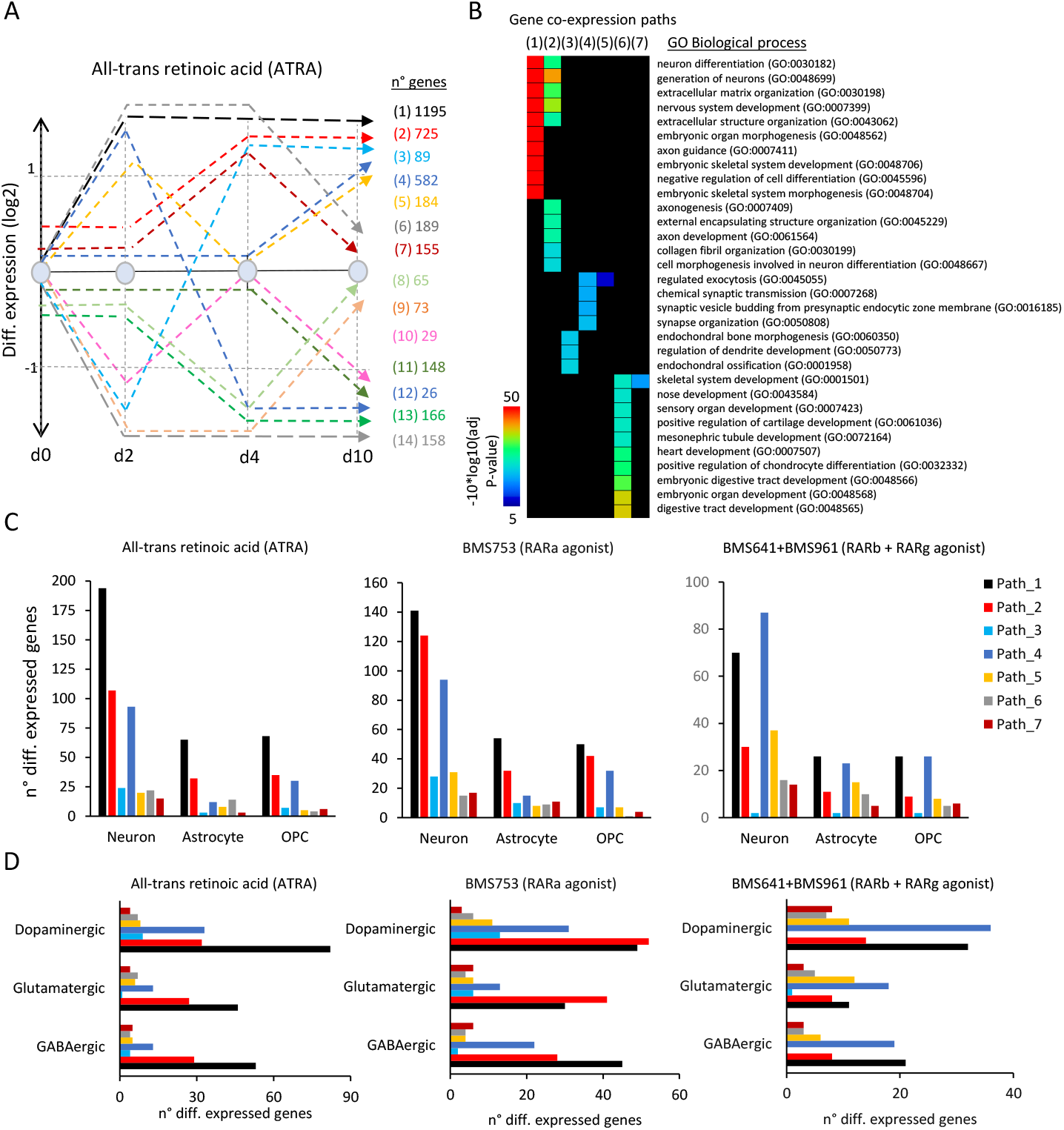
Temporal gene co-expression anaysis during cell specialization driven by retinoids treatment. **(A)** Stratification of the temporal transcriptome profiling during WT P19 cell differentiation driven by ATRA treatment. Transcriptomes were assessed on samples collected at 2, 4 and 10days of treatment. Dashed lines correspond to groupe of differentially co-expressed genes (gene coexpression paths; fold change levels >2). Numbers of genes composing each of the coexpression paths are displayed (right). **(B)** Gene Ontology (GO) analysis on gene co-expression paths displayed in (A) associated to up-regulated events. **(C)** Number of genes per co-expression path corresponding to neuronal, astrocyte of oligodendrocyte precursor (OPC) cell types assessed during ATRA (Left panel), BMS753 (middle panel) and BMS641+BMS961 (right panel) treatment. **(D)** Similar than (C) but corresponding to Dopaminergic, Glutamatergic and GABAergic neuronal subtypes.

As expected, the up-regulated gene co-expression paths of WT P19 cells contained also markers indicative to the various specialized cells, as revealed by comparison with the aforementioned collection resource (**Suppl. Table 1**). As illustrated in **Figure 2C**, 194 genes of the early-responsive gene co-expression Path1 corresponded to neuronal markers, while 65 genes corresponded to astrocyte and other 68 genes corresponded to oligodendrocyte precursors (OPC). The intermediate-responsive Path2 presented 107 genes associated to neurons, 32 to astrocyte, 35 to OPCs, while the late-responsive Path4 presented 93 neuronal markers, 12 genes associated to astrocyte and 30 others associated to OPCs. These kinetics indicate that neuronal differentiation precedes glial cells emergence, in aggreement with previous findings in in-vivo and in-vitro mamalian systems (reviewed in (Miller & Gauthier, 2007; Hirabayashi & Gotoh, 2010)). All other paths presented less than 25 genes corresponding to the aforementioned cells, indicating that early (Path1), middle (path2) and late (path4) responder co-expression paths are the most relevant for describing cell specialization. (**Figure 2C&D**).

While WT P19 cells treated with the RARα agonist BMS753 presented relatively similar transcriptome kinetics; the number of markers associated to specialized cells retrieved on path1 were lower than those observed on the gold-standard ATRA treatment (141 genes associated to neurons, ∼54 genes to astrocyte and 50 genes to OPCs). This observation for Path1 has been further enhanced on cells treated with the RARβ and RARγ agonists (BMS641 & BMS961), including in addition a significant reduction on the number of gene markers associated to specialized cells on path2. In contrary, the number of gene markers observed on the late-responsive path4 remained rather unchanged (87 genes associated to neuronal markers, 23 genes associated to astrocytes and 26 associated to OPCs) (**Figure 2C)**. This observation suggests that while treatment with the combination of the RARβ+γ-agonist gives rise to specialized cells, their differentiation process is delayed over time relative to that observed under the ATRA treatment. This is also supported by the fact that cells under the RARβ+γ-agonist-treatment present gene markers associated to specialized neurons preferentially found on the late-responsive path4 (**Figure 2D**).

### Reconstruction of gene regulatory networks involved in cell specialization driven by retinoid treatment

Our previous work has shown that ligand binding of retinoid receptors triggers a cascade of events which leads to the dynamic activation of other transcription factors (TF) which then regulate their cognate targets. This cascade of transcription regulatory events can be reconstructed by integration of transcription factor-target gene (TF-TG) databases in the temporal transcriptome analysis (Mendoza-Parra *et al*, 2016a; Cahan *et al*, 2014). This way, gene regulatory networks (GRNs) can be reconstructed and master regulator genes deduced (Cholley *et al*, 2018).

Herein we have reconstructed a master GRN from the integration of the temporal transcriptomes assessed on WT P19 cells treated with the pan-agonist ATRA, covering the 10 days of cell treatment, with transcription factor-target gene (TF-TG) annotations (CellNet database: (Cahan *et al*, 2014)). This master GRN, composed by 1,156 nodes (genes) and 17,914 edges (TF-TG relationships), was stratified based on the presence of nodes (genes) associated to a given type of specialized cell type, as described on the aforementioned collection resource (**Suppl. Table 1**) (Tasic *et al*, 2018; Hook *et al*, 2018; Voskuhl *et al*, 2019). (**Figure 3A**). Specifically, the GRN has been first stratified into four major groups: neuronal cell markers (582 nodes; 7,611 edges), astrocytes (161 nodes; 3,007 edges), oligodendrocyte precursors (OPC: 133 nodes; 2,929 edges), as well as a fourth group composed by genes not retrieved on any of the previous classifications (unassigned: 280 nodes; 4,367 edges) (**Figure 3A&B**). Nodes associated to the neuronal group has been further stratified on dopaminergic (214 nodes; 3,457 edges), glutamatergic (111 nodes; 1,568 edges), GABAergic (87 nodes; 1,264 edges) or unassigned neurons (170 nodes; 2,717 edges) (**Figure 3A&C**).

**Figure 3.**
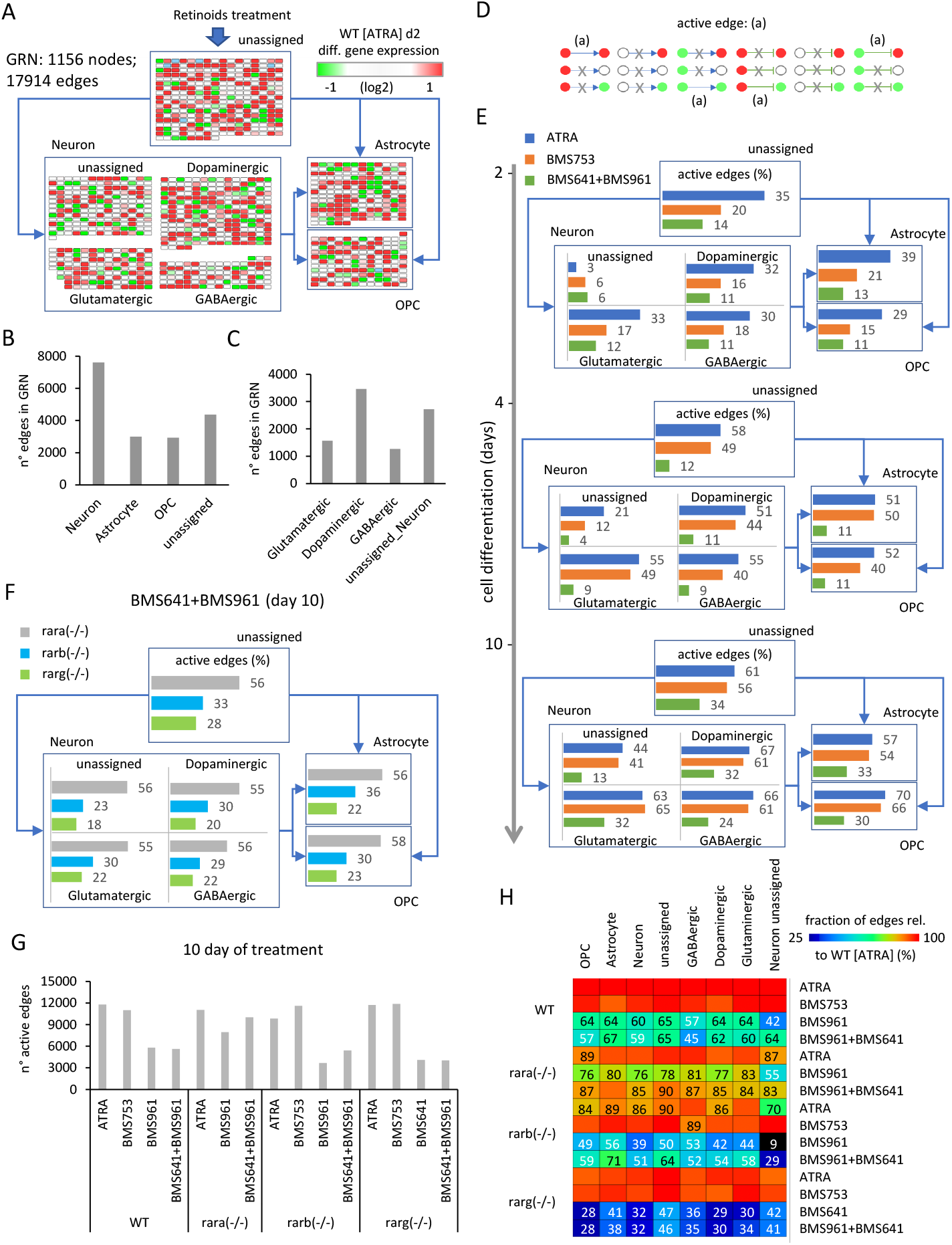
Active gene regulatory wires reconstruction during cell specialization driven by retinoids. **(A)** Structure of the reconstructed gene regulatory network (GRN) displaying differentially expressed genes stratified on four major groups: neuronal cell markers (582 nodes), Astrocytes (161 nodes), oligodendrocyte precursors (OPC: 133 nodes), as well as a fourth group composed by genes not retrieved on none of the previous classifications (unassigned: 280 nodes). Nodes associated to the neuronal group has been further stratified on Dopaminergic (214 nodes), Glutamatergic (111 nodes), GABAergic (87 nodes) or unassigned neurons (170 nodes). For illustration purposes, all edges were removed and replaced by simplified connectors (blue arrows). The color code associated to nodes reflect the differential gene expression levels on WT P19 cells after 2 days of ATRA treatment. **(B)** & **(C)** number of edges interconnecting nodes retrieved on each of the aforementioned groups. **(D)** scheme illustrating all potential types of node’states (active: red, repressed: green, unresponsive:white) and their inter-relationships defined by the illustrated edges (positive regulation: arrow connector, negative regulation: t-shaped connector). “Active edges”, correspond to transcriptionally-relevant nodes/edges relationships, and are conserved during the analytical procressing of the GRN illustrated in (A). **(E)** Temporal transcriptional evolution of the reconstructed GRN during WT P19 cell differentiation. Illustrated barplots correspond to the fraction of active edges (as defined in (D)) relative to the total edges (displayed in (B) & (C)) issued from the treatment with either the all-trans retinoic acid (ATRA), the RARa-specific agonist BMS753 or the combination of the RARb and RARg agonists (BMS641+BMS961). **(F)** barplots corresponding to the fraction of active genes after ten days of treatment with the RARb and RARg agonists (BMS641+BMS961) of P19 RAR-null mutant lines. **(G)** Number of total active edges retrieved on GRNs issued from 10 days of treatment with the indicated retinoids and over the different P19 lines. Notice that the number of active edges on the Rara(-/-) line treated with the combination of the BMS641+BMS961 agonists leads similar levels than those observed on the gold-standard wild-type (WT) line treated with the pan-agonist ATRA. **(H)** Fraction of active edges (relative to the WT line treated with ATRA) associated to markers corresponding to the classification of specialized cells retrieved in (A) relative to those observed on the gold-standard wild-type ATRA treated sample. Fraction levels higher than 90% are only displayed with the heatmap color-code (red).

One of the major advantages of working with reconstructed GRNs is the fact that the relevance of the system can be challenged by the coherence of the interconnected players. In this case, we define as an “active edge”, a set of two nodes being differentially responsive and interconnected with a transcription regulation relationship (active or repressed regulation) coherent with the gene expression status of the interconnected nodes (e.g. active genes requires to be interconnected by active transcription regulation edges; **Figure 3D**). Hence, during differentiation of WT P19 cells treated with the pan-agonist ATRA, the fraction of active edges passes from ∼30% to ∼50% and finally ∼60% when evaluating readouts at 2, 4 and 10 days of treatment associated to specialized cell-types (**Figure 3E**). Interestingly, WT P19 cells treated with the RARα agonist (BMS753) present a lower number of active edges after 2 days of treatment (21% for astrocytes, 15% for OPCs and ∼17% for the specialized neurons), but recovered similar levels than those observed with the ATRA treatment in the late time-points. In contrary, the use of the combination of the RARβ+γ-agonist (BMS641+BMS961) raise the levels of active edges to barely ∼30% after 10 days of treatment (**Figure 3E**). While this poor performance is also observed on P19 lines deficient for the RARβ or the RARγ receptor, the Rara(-/-) P19 mutant line revealed a significant gain on the number of active edges (56% for astrocytes, 58% for OPCs and ∼55% for specialized neurons) (**Figure 3F & Supplementary Fig. S4**).

Indeed, while GRNs corresponding to WT P19 cells treated with BMS961 & BMS641 agonists present ∼half of active edges observed on ATRA treatment conditions (5,628 edges), P19 Rar-alpha (-/-) mutant cells under the same retinoids’ treatment leads to GRNs presenting 10,044 edges interconnecting responsive genes (**Figure 3G**). The comparison of the number of active edges under different conditions with those observed on the gold-standard P19 WT ATRA treatment reveals that the Rara (-/-) mutant treated with the BMS961 & BMS641 agonists leads to a recovery of >80% for all cell specialized groups (**Figure 3H**). In contrary, the same treatment on WT P19 cells reaches levels of ∼60% for most of the groups, with the exception of the GABAergic neuron subtypes were only 45% of edges observed on ATRA treatment are recovered.

Overall, the reconstructed GRN describing cell types specialization during retinoid-driven cell differentiation reveals the fraction of reactivated edges by the synergistic activation of the RARβ and RARγ receptors, notably in the Rara mutant line.

### An enhanced restoration of neuronal cell specialization driven by RARγ and RARβ receptors requires to bypass gene programs controlled by the RARα receptor

A deeper analysis of the reconstructred GRN revealed a 2-fold increase in the number of active edges for P19 Rara(-/-) mutant relative to the WT line when treated with the RARβ+γ agonists (BMS641, BMS961). Such enhanced performance could be explained by a functional redundancy of RAR subtypes, as previously demonstrated during the early phases of endodermal (F9) and neuronal (P19) cell differentiation (Taneja *et al*, 1996, 1995; Roy *et al*, 1995). Specifically, we speculate that, in absence of the RARα receptor, the synergistic activation of the RARβ and RARγ receptors could drive the activation of RARα-specific programs. Similarly, such RARα−specific programs might remain “inhibited” in WT cells (for instance due to the unliganded binding of the RARα receptor) despite the combined exposure to RARβ+γ agonists.

To address this hypothesis, we first identified the RARα agonist (BMS753)-specific programs, corresponding to the common active edges between WT, Rarb(-/-) and Rarg(-/-) mutant lines treated with this synthetic ligand (**Figure 4A**). The obtained 9,328 active edges were then intersected with those observed on WT or Rara(-/-) mutant line treated with the RARβ+γ agonists, to reveal those programs commonly activated by both treatments (3,806 active edges), as well as those specifically activated by the RARα agonist (BMS753) but inhibited in the WT line in despite of the combined exposure to the RARβ+γ agonists (3,830 active edges).

**Figure 4.**
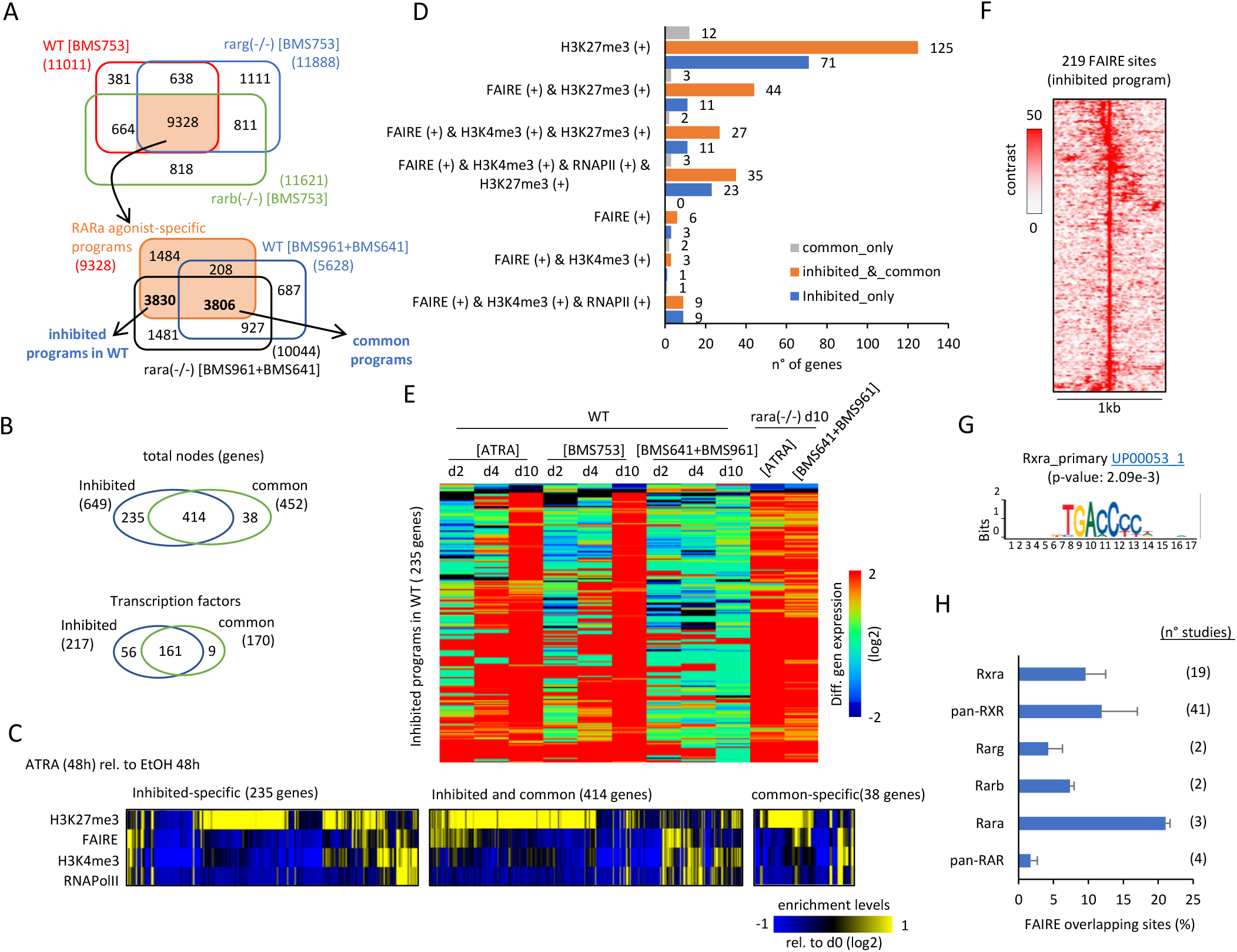
Identification of a subset of active edges remained inhibited by the unliganded RARa receptor in P19 WT cells during the synergistic activation of the RARb and RARg. **(A)** Top panel: Venn-diagram revealing the RARa-specific programs corresponding to the common active edges retrieved in WT, rarg(-/-) and rarb(-/-) P19 lines treated with the RARa agonist BMS753 (9,328 active edges). Bottom panel: Venn-diagram revealing the “common programs” (3,806 active edges) driven by the RARa agonist and those responding to the synergistic activation of the RARb+RARg receptors; as well as a subset of active edges specifically driven by the RARa agonist BMS753 (3,830). This last subset is defined herein as “inhibited programs by the unliganded RARa”, because they remain unresponsive on WT cells treated with the combination of the RARb+RARg ligands (BMS641+BMS961), but they are reactivated on the Rara(-/-) line. **(B)** Top Venn-diagram: comparison between the number of genes retrieved in the “inhibited” and the “common” programs highlighted in (A). Bottom Venn-diagram: comparison between the number of transcription factors retrieved in the “inhibited” and the “common” programs highlighted in (A). **(C)** Heat map illustrating the promoter epigenetics status (repressive mark H3K27me3 and the active mark H3K4me3), the chromatin accessibility (FAIRE), and the transcriptional response (RNA Polymerase II (RNAPII)) of genes specific to the “inhibited” or the “common” program as well as those shared between these two programs after two days of ATRA treatment. **(D)** number of genes presenting the indicated promoter epigenetic combinatorial status in the conditions illustrated in (C). (E) Heat map displaying the differential expression levels for genes associated to the “inhibited”-specific program at different time-points and retinoids treatment. Notice that while most of these genes remained unresponsive when treated with the combination of the RARb+RARg ligands (BMS641+BMS961) in wild-type (WT) cells, they are upregulated on the Rara(-/-) line. **(F)** Open-chromatin FAIRE sites retrieved on the promoters of the “inhibited programs by the unliganded RARa”. (G) Motif analysis performed on the FAIRE sites presented in (F), revealing the enrichment of the Rxra primary motif. (H) Binding sites enrichment analysis performed on the aforementioned FAIRE sites, by comparing with 71 RXR or RAR ChIP-seq publicly available profiles (NGS-QC Generator database: https://ngsqc.org/).

A close look at the “common” and “inhibited” programs revealed that, despite their distinct number of edges, most of the genes composing the “common” program are also part of the “inhibited” program (414 from 452 genes), and this observation is also conserved for the involved transcription factors (161 shared TFs; **Figure 4B**). This observation suggests that gene expression for the “common” and “inhibited” programs are differentially controlled by other molecular factors in addition to TF regulation. To address this hypothesis, we have evaluated their promoter epigenetics status (defined by the repressive mark H3K27me3 and the active mark H3K4me3), their chromatin accessibility (revealed by FAIRE-sequencing assays), and their transcriptional response (revealed by the enrichment of the RNA Polymerase II (RNAPII)) after two days of ATRA treatment (GSE68291; (Mendoza-Parra *et al*, 2016a)). This enrichment analysis (relative to those observed on EtOH vehicle treatment) revealed that genes being either specifically “inhibited” in WT (235 genes), shared between the “inhibited” and the “common” programs (414 genes) or those being specifically-associated to the “common” programs (38 genes) are preferentially repressed at 48h of ATRA treatment, as revealed by the enrichment of the H3K27me3 modification (**Figure 4C & 4D**).

Genes associated to the “inhibited” program are induced between the 4 and 10 days of ATRA treatment, or the synthetic RARα agonist (BMS753); but they remain unresponsive in presence of the combined exposure to the RARβ+γ agonists. This being said, the combined exposure to the RARβ+γ agonists lead to their gene induction on P19 Rara(-/-) mutant line (**Figure 4E**).

With the aim of confirming the role of the RAR-alpha receptor on the predicted “inhibited” program, we have performed an enrichment analysis on their associated 219 FAIRE sites (**Figure 4F**), by comparing them with ChIP-seq binding sites collected from the public domain. Specifically we have used a collection of more than 40 thousand public mouse ChIP-seq datasets, collected as part of our NGS-QC database (https://ngsqc.org/), among which 71 ChIP-seq public profiles correspond to RXR or RAR transcription factors (Blum *et al*, 2020; Mendoza-Parra *et al*, 2016b, 2013). This analysis revealed that ∼24% of the FAIRE sites associated to the “inhibited” program were enriched for RARα binding sites, ∼20% to pan-RXR sites; further supported by a motif analysis revealing the enrichment of the RARα primary motif (**Figure 4G & H**), confirming their transcriptional response driven by the RARα/RXR heterodimer.

With the aim of summarizing this information within a gene regulatory wiring, we have first assembled the FAIRE & H3K27me3 associated “inhibited” programs into a GRN, complemented by edges issued from public ChIP-seq binding sites associated to the various RAR and RXRα receptors, as well as the RXRα primary motif discovery. This summarized “inhibited RXR/RAR” GRN is composed by 85 Nodes and 160 edges, on which each node has been highlighted on the basis of their promoter epigenetic status (**Figure 5**). To further enhance the relevance of master transcription factors within this “inhibited” network, we have computed their master regulatory index by simulating transcription regulation cascades over the complete reconstructed GRN (described on Figure 3A; TETRAMER: (Cholley *et al*, 2018)). A ranking of the TFs on the basis of their master regulatory index allowed to identify a set of 22 transcription factors able to regulate more than 70% of the ATRA-driven gene programs (**Figure 5 A & B**). 15 of them present a transcriptionally active signature after 2 days of treatment, as revealed by their FAIRE-associated promoter status, while the remaining seven are rather repressed (H3K27me3-associated promoter status) (**Figure 5C**). Interestingly this last group of TFs is composed by players like the T-box family member Tbx18 – known to be regulated by retinoic acid during somitogenesis (Sirbu & Duester, 2006); the transcriptional repressors Hic1 – known to have a role in neural differentiation and tumor suppression in the central nervous system (Rood & Leprince, 2013) - and Hes6 - known to promote cortical neuron differentiation through the repression of the TF Hes1 (Gratton *et al*, 2003, 6), the mesoderm specific factor Tcf21, the RNA binding protein Csdc2; the Nuclear factor I-C (Nfic) - known to regulate cell proliferation and differentiation in the central nervous system notably by modulating the expression of the miR-200b (Huang *et al*, 2021), and Zfp827 - also known as ZNF827 - recently shown to negatively regulate neuronal differentiation through the expression of its circular RNA (Hollensen *et al*, 2020). Importantly, all these repressed factors appear interconnected within the reconstructed GRN and preferentially associated to the RARβ or RARγ receptors (**Figure 5D**).

**Figure 5.**
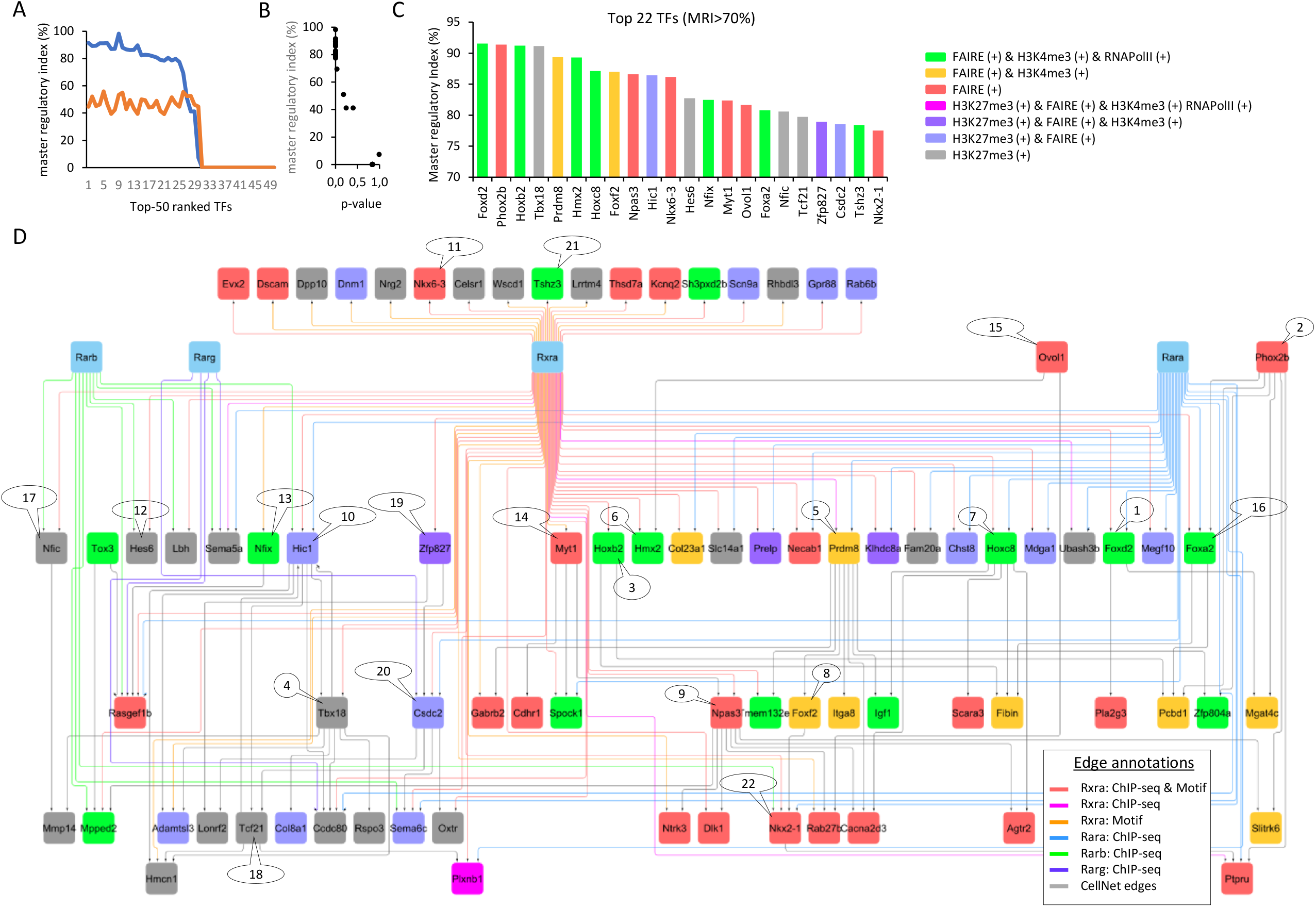
A gene regulatory view of the master players inhibited by the unliganded RARa receptor during neuronal cell specialization. **(A)** Top-50 transcription factors retrieved within the “inhibited program” ranked on the basis of the fraction of downstream controled genes within the reconstructed GRN (Figure 4A) (blue line). The orange line corresponds to the fraction of downstream controled genes predicted on a randomized GRN (Master regulatory index (MRI) computed as described in TETRAMER; Cholley et al…). **(B)** Confidence associated to the TFs’ ranking. Notice that a MRI > 70% present the most confident p-values. **(C)** Transcription factors (22) presenting a MRI>70% and colored on the basis of their promoter epigenetic combinatorial status. **(D)** Gene co-regulatory view of the 22 TFs, illustrating their most relevant (co)regulated players and including their known relationships with RXRa and RAR nuclear receptors.

Among the FAIRE-associated factors, several of them present a RXRα binding site, including the zinc-finger transcription factor Tshz3 – recently described as a “hub” gene involved in early cortical development (Caubit *et al*, 2016) - ; Nfix, recently shown to drive astrocytic maturation within the developing spinal cord (Matuzelski *et al*, 2017); or the Homeodomain transcription factors Hoxb2, Nkx6-3 (involved in development of the central nervous system; the homeobox family factor) or Nkx2-1 – known to control GABAergic interneurons and oligodendrocyte differentiation, and more recently described as driving astroglia production by controlling the expression of the glial fibrillary acidic protein (GFAP) (Minocha *et al*, 2017, 2). Similarly, the homeobox factor Hmx2, the Myelin Transcription Factor 1 (Myt1) or the Neuronal PAS domain protein 3 (Npas3) do present proximal RXRα binding sites; while other players like the Forkhead transcription factors Foxa2, Foxd2 or Foxf2, the homeobox factor Hoxc8, or the histone lysine methyltransferase factor Prdm8 do present in addition a previously-described proximal RARα binding site (**Figure 5D**). Interestingly, Prdm8 appears as a central player within the reconstructed GRN. Indeed, Prdm8 controls 7 other factors known to be highly expressed on nervous tissue: Gabrb2 (the β2 subunit of the GABA_A_ receptors, known to play a crucial role in neurogenesis and synaptogenesis (Barki & Xue, 2022)), the transmembrane protein Tmem132e, Zfp804a (known to regulate neurite outgrowth and involved in neuronal migration (Deans *et al*, 2017)), Igf1 (Insulin-like growth factor 1; synthesized by dopamine neurons (Pristerà *et al*, 2019)), Foxf2 (known to be expressed in neural crest cells leading to pericytes(Reyahi *et al*, 2015, 2)), the Integrin alpha 8 (Itga8) known to regulate the outgrowth of neurites of sensory and motor neurons, and the Calcium Voltage-Gated Channel Auxiliary Subunit Alpha2delta 3(Cacna2d3), known to be essential for proper function of glutamatergic synapses notably on the auditory brainstem (Bracic *et al*, 2022). As a whole, this highlighted Prdm8 regulome appears as a critical player for controlling neuronal differentiation and specialization, in agreement with previous reports on mouse and human differentiation systems (Cypris *et al*, 2020, 8; Inoue *et al*, 2015, 8; Ross *et al*, 2012, 8). Furthermore, our data indicates that Prdm8 as we as all other factors composing the illustrated regulome in **Figure 5**, are driven by the RARα binding sites but can be controlled by the RARβ and RARγ receptors in the absence of the RARα receptor.

## Discussion

How cells respond to different signals to develop along defined cell lineages is a key open question to understand physiological cell differentiation, leading to the formation of organs and tissues, but also events like in-vitro cell reprogramming and even tumorigenesis. In this study, we specifically address the role of retinoids on activating major gene regulatory wires driving neuronal cell lineage and notably cell specialization. Previously we have dissected the major retinoid-driven gene regulatory programs, leading to neuronal precursor formation, notably by evaluating the relevance of the activation of the RARα nuclear receptor in P19 cells with the synthetic agonist BMS753 (Mendoza-Parra *et al*, 2016a).

While we have also showed in our previous study that activation of the RARβ or RARγ nuclear receptors by their cognate BMS641 or BMS961 synthetic agonists is insufficient to promote neuronal differentiation, others reported that in long-term culture conditions which included embryoid body formation, RARγ-specific ligand could induce the formation of GABAergic neurons while RARα induced dopaminergic neurons (Podlesny-Drabiniok et al., 2017). In this study, we addressed neuronal differentiation in long-term culture conditions, but we have kept a monolayer culture strategy because it is known that cell-cell contact interactions retrieved on either 2-dimensional (monolayer) or 3-dimensional cultures could lead to different outcomes, as highlighted by the cellular complexity observed on cerebral organoid cultures (Lancaster *et al*, 2013). We have shown herein that activation of the RARβ receptor does not lead to mature neurons, neither to other specialized cells, while activation of the RARγ nuclear receptors gives rise to lower yields of cell specialization than that observed when using the pan-agonist ATRA or the RARα-specific agonist BMS753. Surprisingly, their synergistic activation gave rise to high yields of maturation, inlcuding specialized neuronal subtypes as well as to other glial cells.

Previous studies demonstrated a redundancy for the activation of certain genes by distinct RXR/RAR receptors, and notably on RAR-null mutant lines, suggesting that in the absence of a given RAR nuclear receptors, the remaining isotypes could compensate for such disfunction (Taneja *et al*, 1996; Roy *et al*, 1995; Chiba *et al*, 1997). Similarly, a synergistic 24h activation by combining RAR isotype agonists has been attempted with P19 embryoid bodies in a recent study, suggesting that the synergistic activation of RARα and RARβ agonists might lead to TH+ dopaminergic neurons, while RARγ and RARβ (or RARγ and RARα) might have a preference to induce Drd2+ neuronal subtypes (Podleśny-Drabiniok *et al*, 2017). Altogether, these studies clearly highlight the redundancy between RAR isotypes, as is further supported with Rar-null mutant experiments illustrated here. Indeed, we clearly demonstrate that Rara KO cells present an enhanced cell specialization yield relative to the wild-type situation. Furthermore, we have decorticated the gene programs that are inhibited by the potential action of the unliganded RARα receptor, notably by observing their activation on the Rara-null line via the synergistic action of the RARβ and RARγ agonists (BMS641+BMS961). Among them we have revealed the inhibition of Prdm8, a member of the family of histone methyltransferases, shown to play a role in the development of brain structures, notably by its capacity to regulate the transition from multipolar to bipolar morphology of cortical neurons (reviewed in (Leszczyński *et al*, 2020)).

In summary, this study provides a systems biology view of the gene programs behind the previously observed redundancy between RAR receptors, paving the way for their potential use for directing cell specialization during nervous tissue formation.

## Materials and Methods

### Cell culture

P19 cells were grown in DMEM supplemented with 1 g/L glucose, 5% FCS, and 5% delipidated FCS. P19 EC cells were cultured in monolayer on gelatin-coated culture plates (0.1%). For cell differentiation assays, all-trans retinoic acid (ATRA) was added to plates to a final concentration of 1 μM for different exposure times. For treatment with RAR subtype-specific agonists, cells were incubated with BMS961 (RARg-specific; 0.1 μM), BMS753 (RARa-specific; 1 μM), and/or BMS641 (RARb-specific; 0.1 μM). After four days of treatment with either of the aforementioned retinoids, medium was replaced by neurobasal medium (ThermoFisher Scientific ref: 21103049) supplemented with N2 (ThermoFisher Scientific ref: 17502048) and B27 devoid of vitaminA (ThermoFisher Scientific ref: 12587010) and cultured for other 6 days.

### Immunohistochemical staining

After 10 days of induced differentiation cells were fixed with 4% paraformaldehyde (Electron Microscopy Sciences), followed by 3 × 5 min washes in PBS. Cells were permeabilized (Triton 0.1% in PBS; 15 min at room temperature) and blocked (10% heat inactivated FCS in PBS) during 1 h at room temperature. Cells were washed 3 × 5 min in permeabilization buffer, then incubated with the primary antibodies antibeta III Tubulin/anti-TUBB3 (Abcam: ab14545) or anti-MAP2 (ab32454). After one hour incubation, cells were washed 3 × 10 min with permeabilization buffer followed by incubation with a secondary antibody (Donkey anti-mouse IgG (H+L) Antibody Alexa 555: Invitrogen A-31570; Donkey anti-rabbit IgG (H+L) antibody Alexa 488; Invitrogen A-21206) and/or DAPI (Invitrogen: D3571). After 1 hour at room temperature, cells were washed for 3 × 10 min in permeabilization buffer, twice with milli Q water and finally mounted on microscope slides.

### RT-qPCR and RNA-sequencing

Total RNA was extracted from P19 cells treated with either ATRA or RAR-specific agonists, using the TRIzol RNA isolation reagent (ThermoFisher Scientific ref: 15596026). One microgram of the extracted RNA was used for reverse transcription (HIGH CAPACITY CDNA RT; Applied Biosystems ref: 4368814). Transcribed cDNA was diluted 5-fold and used for real-time quantitative PCR (QuantiTect SYBR Green; Qiagen ref: 204145). RNA-sequencing libraries were produced with the NEBNext® Ultra™ II RNA Library Prep Kit for Illumina (E7770). Libraries were sequenced within the French National sequencing center; Genoscope (150nts pair-end sequencing; NovaSeq Illumina).

### Primary bioinformatics analyses

Fastq files were qualified with the NGS-QC Generator tool (Mendoza-Parra *et al*, 2016b, 2013). Reads from fastq files were mapped to the mouse mm9 reference genome using Bowtie 2.1.0 under by default parameters. Mapped reads were associated to known genes with featureCounts. RNA-Seq analyses were done with DESeq2 R-package. t-distributed stochastic neighbor embedding (t-SNE) analysis was performed with the R package Rtsne. Heatmap matrices display were generated with MeV 4.9.0. Gene Ontology analyses were performed with DAVID Bioinformatics Resources (https://david.ncifcrf.gov/).

### Collection of gene markers associated to specialized cells

Gene markers associated to neurons (1,352 genes), astrocytes (501 genes), oligodendrocyte precursors (OPC: 501 genes) were collected from the supplementary material (Supplementary dataset_S02) of (Voskuhl *et al*, 2019). Gene markers associated to GABAergic (318 genes) and Glutamatergic (311 genes) neuronal subtypes were collected from the Supplementary Table 9 of (Tasic *et al*, 2018). Gene markers for Dopaminergic (Th+) (513 genes) neuronal subtypes were collected from the Table S2 of (Hook *et al*, 2018). This assembled collection is available in our supplementary material (**Suppl. Table S1**).

### Promoter epigenetic status analysis and RAR/RXR enrichment

P19 epigenetic readouts assessed for the repressive mark H3K27me3 and the active mark H3K4me3, the chromatin accessibility -revealed by FAIRE-sequencing profiling- and the transcriptional response (RNA Polymerase II) after two days of ATRA treatment were collected from our previous published study (GSE68291; (Mendoza-Parra et al, 2016a)). Normalized enrichment levels at gene promoter regions, relative to EtOH control profiles, were used for predicting their epigenetic combinatorial status. Enrichment heat maps and mean density plots for FAIRE readouts at gene promoter regions (+/- 500 bp) were obtained with seqMINER 1.3.4.

FAIRE sites motif analysis has been performed with the MEME Suite 5.4.1. RXR/RAR enrichment on FAIRE sites were inferred by comparing them with > 40 thousand mouse ChIP-seq binding sites collected from the public domain, as part of our NGS-QC database (https://ngsqc.org/). Among all mouse ChIP-seq collected data, 71 ChIP-seq public profiles correspond to RXR or RAR transcription factors(Blum *et al*, 2020; Mendoza-Parra *et al*, 2016b, 2013).

### Gene regulatory networks reconstruction

Temporal transcriptomes issued from ATRA treatment were integrated with a collection of Transcription factor-target gene relationships (CellNet; (Cahan *et al*, 2014)) with the help of our previously developed Cytoscape App, TETRAMER (Cholley *et al*, 2018). TETRAMER has been also used for identifying the top-22 master transcription factors (Master regulatory Index > 70%). Gene co-regulatory wires visualization has been performed with Cytoscape 3.8.2.

## Data access

All RNA-sequencing datasets generated on this study have been submitted to the NCBI Gene Expression Omnibus (GEO; http://www.ncbi.nlm.nih.gov/geo/) under accession number GSE204816.

## Acknowledgements

We thank all the members of our laboratory and the Genoscope sequencing platform for discussion. This work was supported by the institutional bodies CEA, CNRS, and Université d’Evry-Val d’Essonne. E.M. was supported by Genopole Thematic Incentive Actions funding (ATIGE-2017); A.K. by the ‘‘Fondation pour la Recherche Medicale’’ (FRM; funding ALZ-201912009904); F.S. by the funding 2OI9-L22 from the Institut National du Cancer (INCa) and A.B. by the funding 2020-181 (INCa).

## Author contributions

**Elodie Mathieux & Aysis Koshy:** Investigation ; Methodology ; Formal Analysis. **François Stüder :** Data curation; Software; Formal analysis. **Aude Bramoulle :** Resources; Methodology. **Michele Lieb:** Resources; Methodology. **Bruno Maria Colombo:** Writing – review & editing. **Hinrich Gronemeyer :** Supervision; Funding acquisition; Writing – review & editing. **Marco Antonio Mendoza-Parra:** Conceptualization; Formal analysis; Supervision; Funding acquisition; Writing-original draft.

## Conflict of interest

The authors declare that they have no conflict of interest.

## Notes

### Competing Interest Statement

The authors have declared no competing interest.

### Summary of Updates

The title has been enhanced.

