## Supplementary Figures for "Synergistic activation of RARb and RARg nuclear receptors restores cell specialization during stem cell differentiation by hijacking RARa-controlled programs"

### SUPPLEMENTARY MATERIAL

#### Synergistic activation of RARb and RARg nuclear receptors restores cell specialization during stem cell differentiation by hijacking RARa-controlled programs

**Aysis Koshy [1,*], Elodie Mathieux [1,*], François Stüder [1], Aude Bramoulle [1], Michele Lieb [2], Bruno Maria Colombo [1], Hinrich Gronemeyer [2] and Marco Antonio Mendoza-Parra [1,3]**

**Supplementary Figure S1.** Generation and validation of P19 cells deficient for each of the RAR receptors.

**Supplementary Figure S2.** Differential expression response of the neuronal precursor gene Tubb3, the astrocyte-related marker Gfap, as well as the oligodendrocyte marker Olig2 in the context of the P19 cells deficient for each of the RAR receptors.

**Supplementary Figure S3.** Number of differentially expressed genes observed on global transcriptomes and their comparison with genes associated to specialized cells.

**Supplementary Figure S4.** Gene regulatory network (GRN) displaying the differential gene expression response during the 10 days of treatment.

**Supplementary Table S1.** Collection of gene markers associated to specialized cells.

**Supplementary Table S2.** Globla differentially expression levels relative to Ethanol control samples.

A

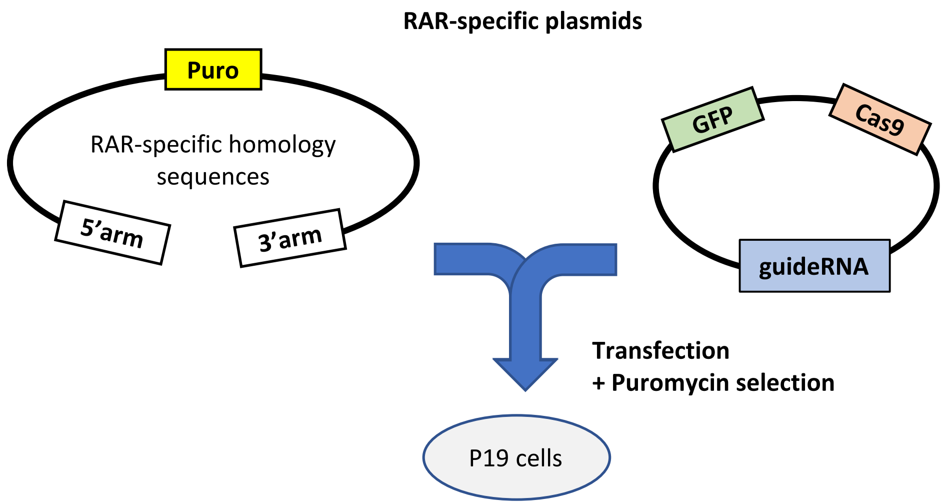


B

| Plasmid name | Santa Cruz Biotech Ref. |
|---|---|
| Plasmid CRISPR/Cas9 KO RARγ/Retinoic Acid Receptor γ (m)<br>Plasmid HDR RARγ/Retinoic Acid Receptor γ (m) | sc-422600<br>sc-422600-HDR |
| Plasmid CRISPR/Cas9 KO RARβ/Retinoic Acid Receptor β (m)<br>Plasmid HDR RARβ/Retinoic Acid Receptor β (m) | sc-432322<br>sc-432322-HDR |
| Plasmid CRISPR/Cas9 KO RARγ/Retinoic Acid Receptor γ (m)<br>Plasmid HDR RARγ/Retinoic Acid Receptor γ (m) | sc-422601<br>sc-422601-HDR |

C

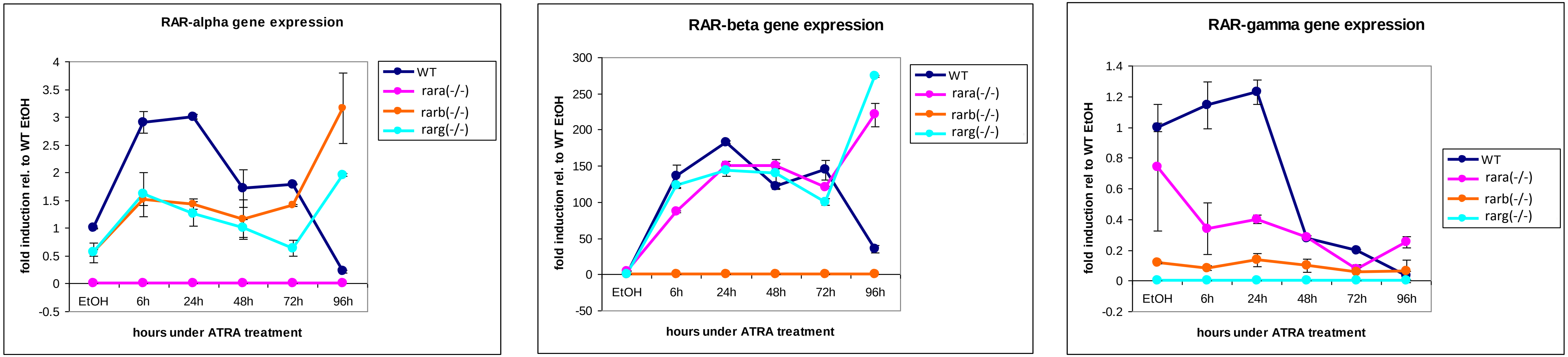


**Supplementary Figure S1. Generation and validation of P19 cells deficient for each of the RAR receptors**. (A) Gene knockout strategy based on targeted CRISPR/Cas9 DNA cleavage and the incorporation of a puromycin resistance cassette by homologous recombination. (B) Reference of the used plasmids, purchased from Santa Cruz Biotechnology. (C) RT-qPCR assay evaluating the expression of each of the RAR receptors during the first 96 hours of P19 cell differentiation driven by All-trans retinoic acid (ATRA) treatment in either WT control cells or the generated rar null mutant lines.

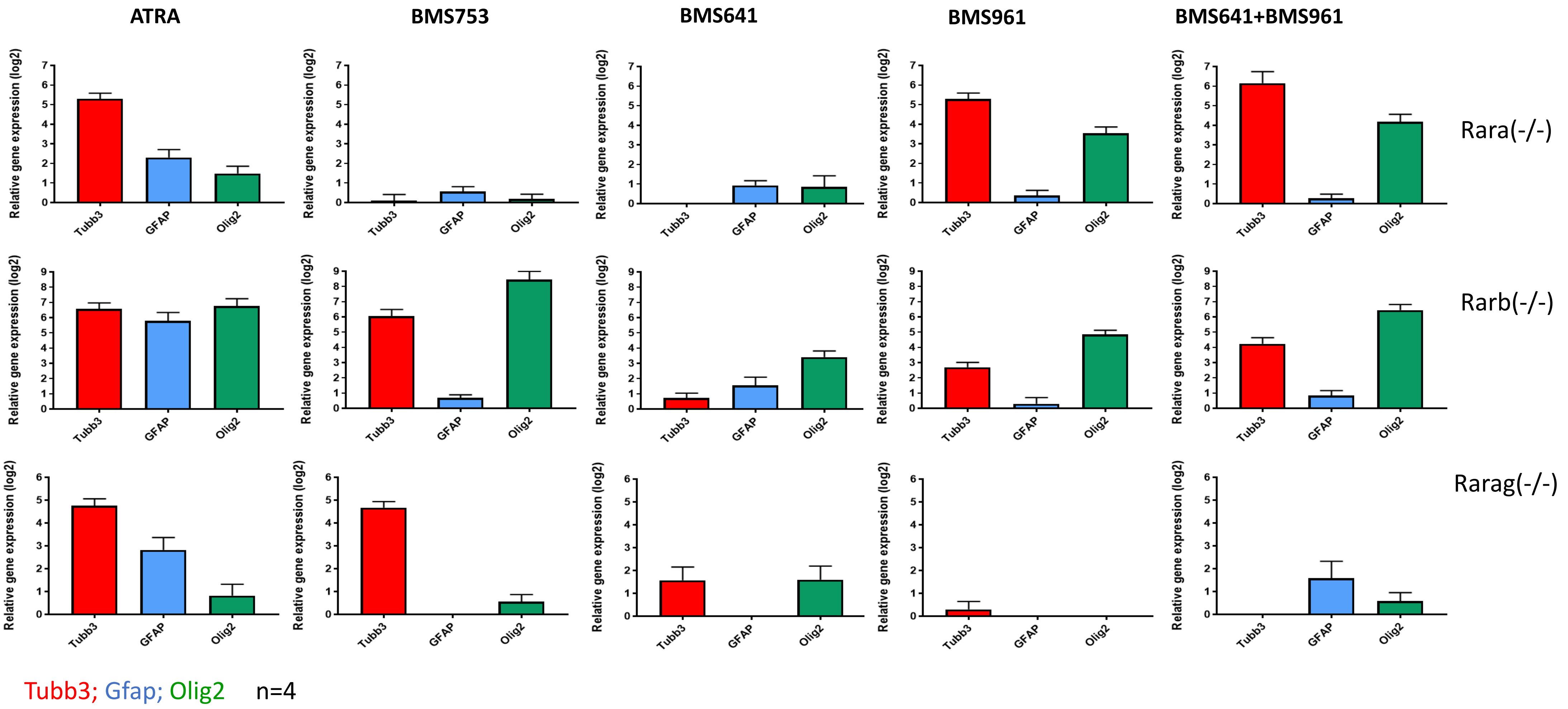


**Supplementary Figure S2. Differential expression response of the neuronal precursor gene Tubb3, the astrocyte-related marker Gfap, as well as the oligodendrocyte marker Olig2 in the context of the P19 cells deficient for each of the RAR receptors.** P19 rar mutant lines were treated (10 days) with either the pan retinoic acid agonist ATRA, or the synthetic agonists BMS753 (Rara-specific), BMS641 (Rarb-specific), BMS961 (Rarg-specific). Illustrated RT-qPCR assays correspond to the average of four independent experiments.

A

| P19 line | treatment | time-point (Days) | DEGs |
|---|---|---|---|
| WT | ATRA | D2 | 1580 |
| WT | ATRA | D4 | 1921 |
| WT | ATRA | D10 | 2483 |
| WT | BMS753 | D2 | 1270 |
| WT | BMS753 | D4 | 1795 |
| WT | BMS753 | D10 | 2423 |
| WT | BMS961+BMS641 | D2 | 1224 |
| WT | BMS961+BMS641 | D4 | 1149 |
| WT | BMS961+BMS641 | D10 | 1718 |
| WT | BMS961 | D10 | 1690 |

| P19 line | treatment | time-point (Days) | DEGs |
|---|---|---|---|
| rara(-/-) | ATRA | D10 | 2333 |
| rara(-/-) | BMS961 | D10 | 1946 |
| rara(-/-) | BMS961+BMS641 | D10 | 2006 |
| rarb(-/-) | ATRA | D10 | 2509 |
| rarb(-/-) | BMS753 | D10 | 2476 |
| rarb(-/-) | BMS961+BMS641 | D10 | 1618 |
| rarg(-/-) | ATRA | D10 | 2536 |
| rarg(-/-) | BMS753 | D10 | 2353 |
| rarg(-/-) | BMS641 | D10 | 1660 |
| rarg(-/-) | BMS961+BMS641 | D10 | 1688 |
| rarb(-/-) | BMS961 | D10 | 1635 |

B

| | Total cell markers* | 1352 | 501 | 501 | 318 | 311 | 513 |
|---|---|---|---|---|---|---|---|
| | Celltypes: | | | | Neuron Subtype | | |
| ID (10 days of treatment) | total induced genes | Neuron | Astrocyte | Oligodendrocyte precursor | GABAergic | Glutamatergic | Dopaminergic |
| WT [ATRA] | 2158 | 401 (29/18) | 100 (19/4) | 131 (26/6) | 92 (28/22) | 78 (25/19) | 150 (29/37) |
| WT [BMS753] | 2128 | 404 (29/18) | 103 (20/4) | 129 (25/6) | 95 (29/23) | 91 (29/22) | 145 (28/35) |
| WT [BMS961] | 1464 | 185 (13/12) | 65 (12/4) | 62 (12/4) | 47 (14/25) | 49 (15/26) | 69 (13/37) |
| WT [BMS961+BMS641] | 1377 | 203 (15/14) | 61 (12/4) | 58 (11/4) | 44 (13/21) | 49 (15/24) | 84 (16/41) |
| rara(-/-) [ATRA] | 2061 | 348 (25/16) | 104 (20/5) | 109 (21/5) | 96 (30/27) | 80 (25/22) | 128 (24/36) |
| rara(-/-) [BMS961] | 1700 | 261 (19/15) | 79 (15/4) | 92 (18/5) | 69 (21/26) | 65 (20/24) | 100 (19/38) |
| rara(-/-) [BMS961+BMS641] | 1831 | 288 (21/15) | 93 (18/5) | 92 (18/5) | 77 (24/26) | 72 (23/25) | 110 (21/38) |
| rarb(-/-) [ATRA] | 2159 | 363 (26/16) | 110 (21/5) | 119 (23/5) | 97 (30/26) | 81 (26/22) | 140 (27/38) |
| rarb(-/-) [BMS753] | 2177 | 387 (28/17) | 112 (22/5) | 130 (25/5) | 93 (29/24) | 77 (24/19) | 138 (26/35) |
| rarb(-/-) [BMS961] | 1341 | 132 (9/9) | 56 (11/4) | 47 (9/3) | 37 (11/28) | 30 (9/22) | 52 (10/39) |
| rarb(-/-) [BMS961+BMS641] | 1393 | 152 (11/10) | 66 (13/4) | 53 (10/3) | 44 (13/28) | 33 (10/21) | 57 (11/37) |
| rarg(-/-) [ATRA] | 2246 | 406 (30/18) | 109 (21/4) | 127 (25/5) | 100 (31/24) | 83 (26/20) | 149 (29/36) |
| rarg(-/-) [BMS753] | 2101 | 368 (27/17) | 108 (21/5) | 113 (22/5) | 90 (28/24) | 78 (25/21) | 137 (26/37) |

**n°genes (% rel to n° markers / % rel to n° induced genes)**

**Supplementary Figure S3. Number of differentially expressed genes observed on global transcriptomes and their comparison with genes associated to specialized cells. (A)** Differentially expressed genes (DEGs rel to Ethanol-treated control sample) in the WT and RAR mutant P19 lines and under the listed treatments and at different stages (time-points in days) were computed for a log2 fold-change threshold of 2. **(B)** Number of Induced genes in the listed conditions (ID) after 10 days of treatment, and the fraction corresponding to Neuron, Astrocyte or Oligodendrocyte precursor cell types, as well as to the indicated neuron subtypes. Fractions (in percent) were calculated relative to either the described total number of genes per cell markers listed on Supplementary Table S1 (*), or relative to the total number of induced genes. Highlighted rows correspond to samples treated with the RAR-beta+RAR-gamma ligands.

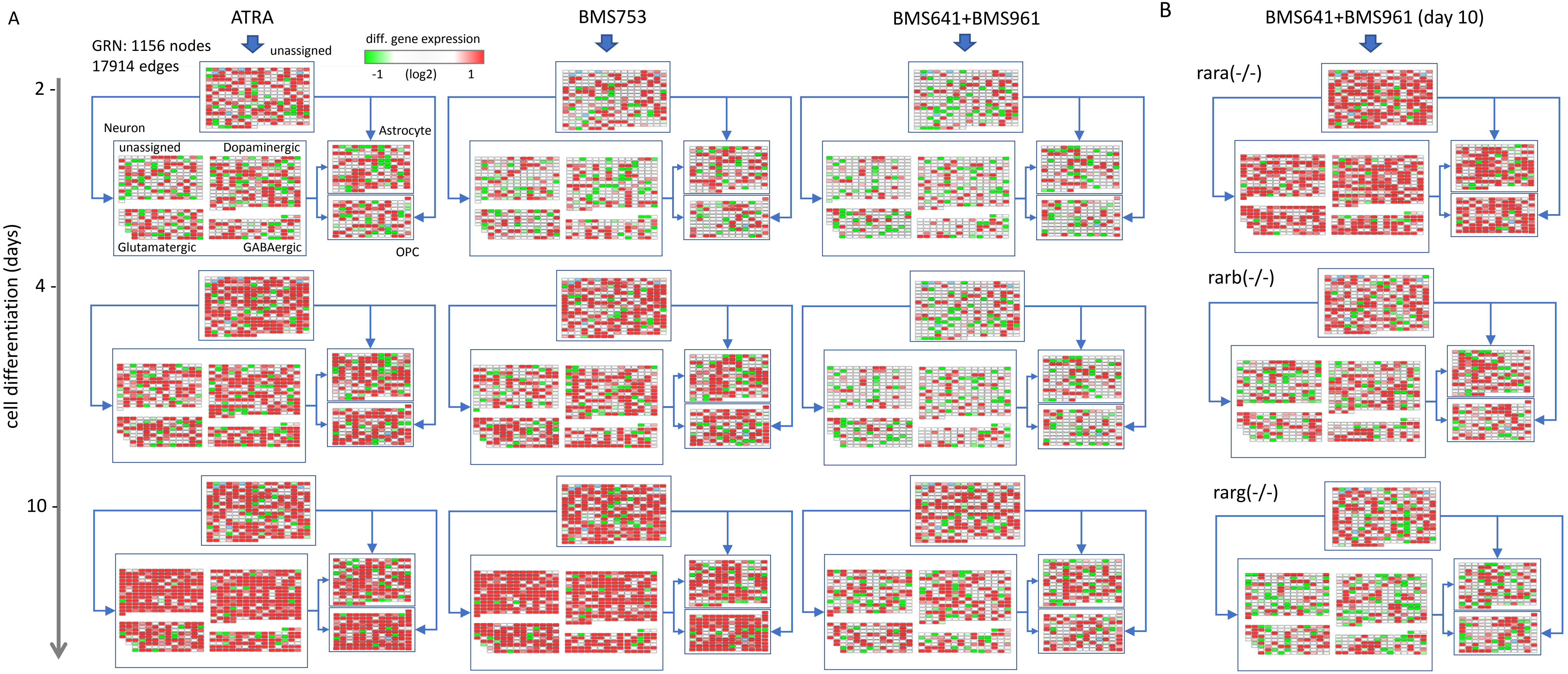


**Supplementary Figure S4. Gene regulatory network (GRN) displaying the differential gene expression response during the 10 days of treatment. (A)** Temporal gene expression response on WT P19 cells driven by the treatment with either ATRA, the RARalpha-agonist BMS753 or the combination of the RARb+RARg agonists (BMS961+BMS641) and visualized within a reconstructed GRN. GRN is composed by 1156 nodes (genes) and 17914 relationships (edges) collected from transcription factor-target gene databases (Cholley et al; 2018). Furthermore, Nodes composing the GRN are stratified into four major groups based on their correspondence to following cell specialization markers: Neurons, Astrocytes, Oligodendrocyte precursors (OPCs) and Unassigned nodes when no correspondence to such markers is retrieved. Nodes associated to the class “Neuron” are further stratified on either Dopaminergic, Glutamatergic, GABAergic or unassigned-neurons. Nodes are colored on the basis of their differential response at the three assessed timepoints. (B) Differential gene expression response observed after 10 days of treatment with the combination of the RARb+RARg agonists (BMS961+BMS641) on the indicated Rar mutant lines.
